# Common Neural Choice Signals reflect Accumulated Evidence, not Confidence

**DOI:** 10.1101/2025.02.13.638046

**Authors:** Kobe Desender, Andi Smet, Deniz Erdil, Esna Mualla Gunay, Yvonne F Visser, Pierre Le Denmat, Hélène Van Marcke

## Abstract

Centro-parietal EEG signals (CPP and Pe) correlate with the reported level of confidence. According to recent computational work these signals reflect evidence which feeds into the computation of confidence, not directly confidence. To test this prediction, we causally manipulated prior beliefs to selectively affect confidence, while leaving objective task performance unaffected. Behaviorally, we found that manipulating prior beliefs causally affected confidence without corresponding changes in accuracy and a negligible effect on reaction times. The EEG data showed a monotonic relation between the reported level of confidence and both CPP and Pe amplitudes. Importantly, this finding is compatible both with the theory that these signals track confidence as well as with the alternative theory that they track accumulated evidence. Critically, both neural signals were insensitive to the influence of prior beliefs on confidence, showing that they reflect the accumulated evidence that is used by the system to compute confidence, not directly confidence. Likewise, oscillatory activity in alpha and beta band was insensitive to the influence of prior beliefs on confidence. Decoding analyses revealed that the brain does hold shared representations for prior beliefs and confidence, and we identified a frontal signal that is sensitive to both confidence and prior beliefs.

## Introduction

Much neuroscientific work unravelling the temporal unfolding of decision confidence has been stimulated by computational work couched within decision making models. One of the most influential accounts of decision making posits that choices are formed based on the accumulation of noisy evidence over time, which continues until the accumulated evidence reaches a predefined boundary (Ratcliff & McKoon, 2008; Tuerlinckx et al., 2001). Several influential papers have demonstrated that confidence can be accounted for within this framework by assuming that it reflects the probability of a choice being correct given the level of accumulated evidence and the time spent accumulating (Kiani et al., 2014; Kiani & Shadlen, 2009). Such models describe the computation of confidence as something that happens at the same time of the decision (i.e. decisional locus) and make the key prediction that neural signatures of confidence should be visible as a decision unfolds. Using EEG recordings in humans, it has been demonstrated that a centro-parietal positivity (CPP) component implements this accumulation-to-bound process (Kelly & O’Connell, 2013; O’Connell et al., 2012). Critical for the current purpose, several studies have demonstrated modulations of the CPP as a function of confidence (Boldt & Yeung, 2015; Desender et al., 2019; Dou et al., 2024; Feuerriegel et al., 2022; Gherman & Philiastides, 2015, 2018; Kopcanova et al., 2023; Lim et al., 2020; Rausch et al., 2020; Stone et al., 2024). The CPP reaches a higher amplitude when people report higher confidence for their decisions. Another set of computational models has described confidence as a post-decision accumulation process (Herregods et al., 2023; Moran et al., 2015; Pleskac & Busemeyer, 2010). In these models, the accumulation process is allowed to continue after crossing the boundary, and the post-decision evidence accumulation informs the computation of confidence. In line with predictions from such a framework, several studies have found that a post-decision centro-parietal EEG signal, often referred to as Error Positivity (Pe), correlates with the level of decision confidence (Boldt & Yeung, 2015; Desender et al., 2019) and changes of mind (Stone et al., 2024), and thus this signal is thought to represent post-decision accumulation (Desender et al., 2021). Note, however, that some investigations have not found the modulation by confidence (Feuerriegel et al., 2022; Ko et al., 2024) or only when presenting post-decision evidence to participants (Grogan et al., 2023).

Recent empirical and computational work has shown that confidence is computed by combining prior beliefs about the ability to perform the task with the accumulated evidence on a single decision (Van Marcke et al., 2024). At the behavioral level, this work showed that exposing participants to overly positive vs. negative feedback or training them on an easy vs. difficult version of a task led them to believe they were very good vs. bad at this task, which in turn causally induced over- vs. under confidence. This effect of prior beliefs was selective for confidence, as it left task performance unaffected. Most important for the current purpose is that this finding could be explained by a computational model in which the accumulation of evidence was unaffected by prior beliefs (i.e. as observed for choices and RTs), and prior beliefs instead selectively altered the probability map used to quantify confidence. That is, the model quantifies confidence as the probability of being correct conditional on evidence and time – and this mapping depends on the prior beliefs. Critically, this model makes a key prediction: common neural choice signals (such as the CPP and Pe) that are often associated with confidence are predicted to instead reflect evidence that is used to compute confidence, but these signals do not directly reflect confidence. Since previous work did not explicitly manipulate prior beliefs, it has not been possible to distinguish between these two possibilities. Therefore, the current work tests the novel prediction that common neural choice signals reflect evidence rather than confidence.

## Methods

### Open Science Statement

All raw data, analysis code and experiment code are openly available at [insert link at publication].

### Participants

The study consisted of a total of 32 Dutch-speaking participants (8 Male, M_age_ = 19.4, SD_age_ = 2.2, range: 18-25) having normal or corrected-to-normal vision. Participants were not allowed to participate if one of the following exclusion criteria was met: having a traumatic brain injury or loss of consciousness in the past years, being pregnant, being under the influence of alcohol or drugs, using a psychotropic medication, having a neurological or psychiatric diagnosis, or having an epileptic disorder. Data from one participant was excluded because they did not complete the experiment, and data from another participant was excluded because of chance level performance. One participant gave the same confidence level on every trial during the orientation task but gave normal responses in all the other tasks. Therefore, only the orientation task was excluded from the analysis. The final sample thus included 30 participants. All participants read and signed the informed consent form approved by the KU Leuven ethics committee before the start of the experiment (code: G-2020-2609-R3(AMD)).

### Task and procedure

The experiment was fully within-subjects and comprised four different phases. In each phase, participants performed a different perceptual decision-making task (i.e. either a color discrimination task, a shape discrimination task, a letter discrimination task, or a line orientation task). In each of these tasks, the stimuli comprised eighty elements (i.e. 80 red or blue dots, 80 squares or circles, 80 x’s or o’s, 80 vertical, or horizontal lines) and the ratio between the number of dominant versus non-dominant elements determined trial difficulty. Theoretically, the most difficult stimulus would thus be 41 vs 39 whereas the easiest stimulus would be 80 vs 0. Each task featured three levels of difficulty which were set beforehand based on our previous studies (Van Marcke et al., 2024) for the color discrimination task (easy: 28 vs 52; medium: 34 vs 46; hard: 37 vs 43), the shape discrimination task (easy: 19 vs 61; medium: 24 vs 56; hard: 32 vs 48), and the letter discrimination task (easy: 17 vs 63; medium: 24 vs 56; hard: 32 vs 48), or established during a pilot study for the line orientation task (easy: 23 vs 57; medium: 24 vs 56; hard: 35 vs 45).

Each of the four different phases of the experiment consisted of a training phase followed by a testing phase. Each training phase comprised 5 blocks of 18 trials and aimed to manipulate the prior belief about the ability to perform the task. There were four different conditions, two of which served to install the prior belief that one is very good at performing the task at hand, and two installed the prior belief that one is very bad at performing the task at hand. Specifically, (i) In the positive feedback condition participants received positive comparative feedback (e.g. you are among the best 90%) every 18 training trials (for a total of 5 times). One out of five times the feedback screen showed ‘moderate’ feedback (e.g. you are at the best 40%) in order to make the feedback appear more realistic. Note that all three difficulty levels were used during training. (ii) In the negative feedback condition, everything was the same as (i) except that participants received negative feedback (e.g. you are among the worst 10%). (iii) In the hard training difficulty condition all training trials were difficult (i.e. aiming for 65% accuracy) and participants received veridical feedback about their performance after each decision (i.e. “correct” or “wrong”). (iv) In the easy training difficulty condition everything was the same as in (iii) except that all training trials were easy (i.e. aiming for 95% accuracy). Note that the positive feedback condition and the easy training condition both induced the belief that participants were good at a task and are henceforth referred to as the positive prior beliefs condition, whereas the other two conditions are referred to as the negative prior beliefs condition. The order of the tasks and manipulations was counterbalanced between participants following a Latin square.

On each trial, a fixation cross was presented for 750ms, the stimulus was presented for 200ms, followed by a blank screen which lasted until participants provided a response by pressing ‘c’ or ‘n’ with the left and right thumb, respectively. After each training phase, participants took part in a testing phase which lasted 3 blocks of 60 trials. Testing phases presented trials of all three difficulty levels. Participants did not receive any feedback about their performance, instead they rated their confidence for each trial using a 6-point scale, with the labels “certainly wrong”, “probably wrong”, “guess wrong”, “guess correct”, “probably correct”, and “certainly correct”. Participants rated their confidence using the ring, middle, and index fingers of both hands by pressing 1, 2, 3, 8, 9, or 0 respectively, at the top of their keyboard. To avoid visual artefacts, the confidence labels were presented throughout the testing phase on top of the screen in grey. After participants indicated their confidence, the corresponding label turned white for 750ms (i.e. to inform that the response was registered). A visual depiction of an example trial is shown in Figure 1A.

**Figure 1.**
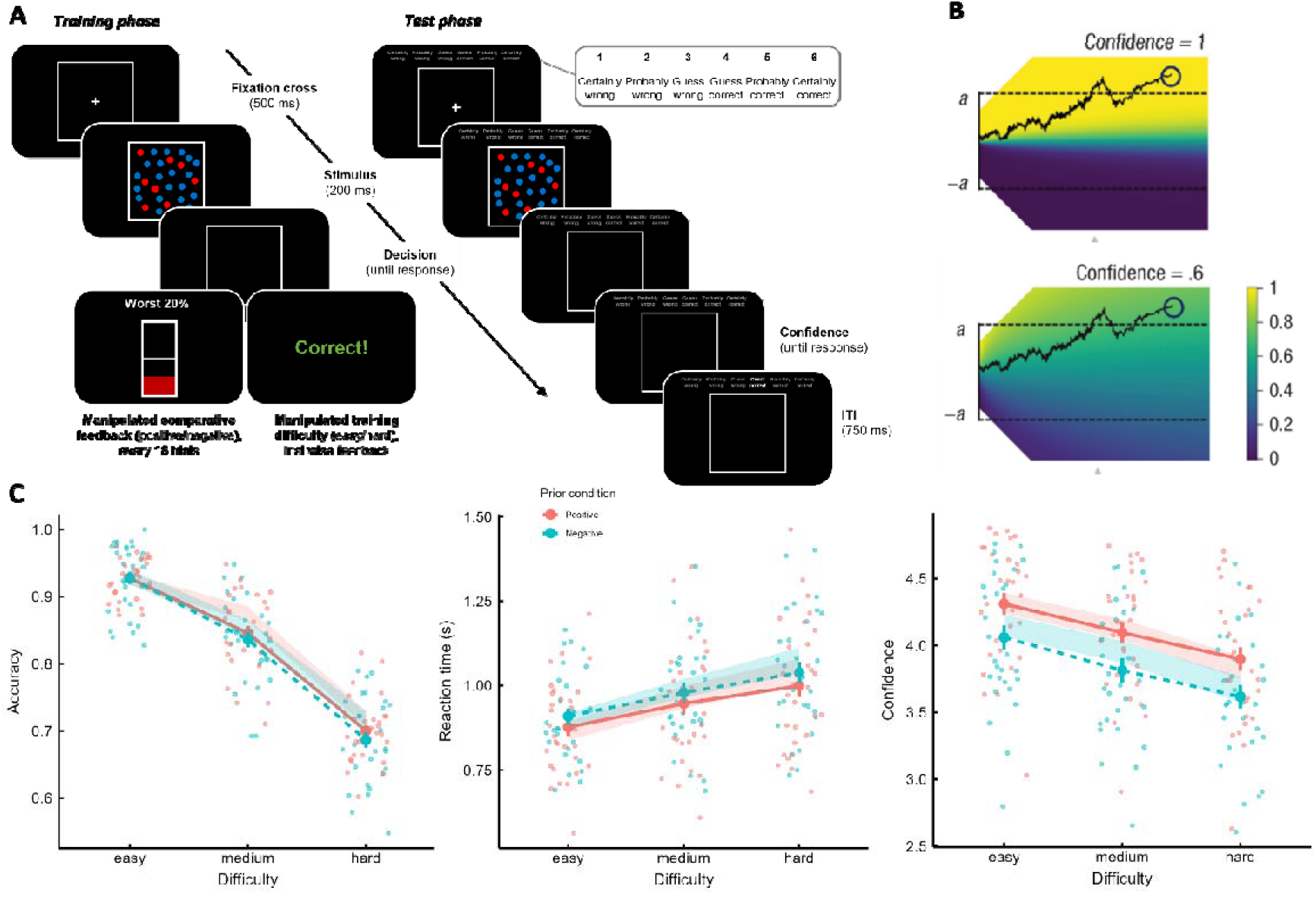
Design and behavioral results. **A.** On each trial, participants made a speeded perceptual decision about a stimulus that was shown for 200ms. There were four different perceptual tasks (only one shown for illustration). During the training phase of each task, participants were told every 18 trials that they performed better compared to a reference group (positive feedback condition), that they performed worse compared to a reference group (negative feedback condition), or they received trial-by-trial feedback while being trained on an easy version of the task (easy training) or a very difficult version of the task (hard training). Each of these manipulations installed the prior belief that participants were very good (i.e. positive prior belief) or very bad (i.e. negative prior belief) at the task, respectively. During the following test phase, participants no longer received feedback but instead rated their confidence on each trial on a six-point scale. **B.** When holding the prior belief that a task is very easy or very difficult, the normative strategy is to report higher/lower confidence even for the exact same trial, because the probability of being correct is effectively higher in the former. This effect of the prior beliefs can be seen in the figure as the difference between the two heatmaps representing the probability correct in both figures. C. Both accuracy and RTs were sensitive to task difficulty, but not to the prior belief conditions. On the contrary, decision confidence strongly depended on the prior beliefs condition: participants consistently reported higher confidence when they believed they were very good versus very bad at the task. As shown by the shaded lines, this pattern was captured by the computational model outlined in B. Note: dots show means of individual participants, large dots show group averages. Error bars show the standard error of the mean across participants, shaded lines show fits of a drift diffusion model including a subjective drift rate parameter controlling for the influence of prior beliefs on confidence. The width of the shaded lines reflects the standard error of the mean of the model fits.

### Behavioral analysis

Only the data of the test phases (i.e. after the prior beliefs were installed) were analyzed. To allow a direct comparison between behavioral and EEG, analysis of behavioral data was done on the EEG preprocessed data, resulting in 5.2% data loss due to EEG preprocessing. In addition, trials with decision RTs slower than 3s were excluded (0.9% data loss) and trials with confidence RTs slower than 3s were excluded (0.3% data loss). Due to the subjective nature of the confidence reports, there was very little data for the lowest two confidence levels. For trials judged as “certainly wrong”, there were only 114 (in the negative condition) and 89 (in the positive condition) datapoints across participants (with 65 of those trials coming from a single participant). For trials judged as “probably wrong”, there were only 217 (in the negative condition) and 133 (in the positive condition) datapoints across participants (with 84 of those trials coming from a single participant). Therefore, we merged the lowest two levels of confidence into a single category and afterwards scaled confidence between 1 and 5. This procedure ensured that there were on average more than 18 trials per confidence level per participant (M_certainly/probably wrong_ = 18.4, M_guess wrong_ = 33.0, M_guess correct_ = 157.8, M_probably correct_ = 258.4, M_certainly correct_ = 201.2). Behavioral data were analyzed in R using mixed-effects models as implemented in the lme4 package (Bates et al., 2015). Model selection was performed by first establishing the fixed effects based on the hypothesis and a random intercept for each participant. Random slopes were included in the model if this increased model fit, as assessed by model comparison using likelihood ratio. Confidence and reaction times were analyzed using linear mixed-effect models, while accuracy was analyzed with general linear mixed-effect models. In case of theoretically relevant null effects, the models were refitted using the BayesFactor package (Morey & Rouder, 2014) in order to acquire Bayes Factors (BFs). Significance testing and model comparison was done using an alpha level of .05. For linear mixed-effect models we report F-values with Satterthwaite’s degrees of freedom, for generalized linear mixed-effect models we report chi square statistics.

### Model fitting

We fit the behavioral data using the same computational model described in Van Marcke and colleagues (2024), which is performed in two successive steps. In the first step, we fit the drift diffusion model (DDM) parameters to the accuracy and reaction time data (Tuerlinckx et al., 2001; Vandekerckhove & Tuerlinckx, 2007). We used quantile optimization to estimate the following five DDM parameters separately for each participant: boundary, non-decision time and three drift rates (one for each difficulty level). In the second step, we fit subjective drift rates to confidence ratings, separately for each participant and condition. The subjective drift rate directly controls the shape of the probability map from which confidence is quantified based on the combination of evidence and RT. Thus, for the same combination of evidence and RT, a higher subjective drift rate will result in higher confidence compared to a lower subjective drift rate. Because model-predicted confidence is given as a probability of being correct (i.e. between 0 and 1), we applied equal-width binning to transform this probability into a 1-5 rating. To account for individual differences and biases in mapping confidence to a categorical scale, we additionally estimated a single bias parameter for each participant. Importantly, the bias parameter was fixed across conditions so it could not explain the prior beliefs effect. The objective function to minimize for this second step was the mean square difference between the proportion of observed vs. predicted confidence ratings for each confidence level. The optimization for both steps was done using a differential evolution algorithm, as implemented in the DEoptim R package (Mullen et al., 2011). Population size was set to ten times the number of parameters to estimate. Optimization continued until no improvement in the objective function was observed for the last 100 generations or until reaching 1000 generations (the latter stopping criterion was never met).

### EEG Recording and preprocessing

Participants sat in a dimly lit, electrically shielded room. EEG data were recorded using a Biosemi system with 64 electrodes. The data were continuously recorded using BioSemi ActiveTwo amplifiers, sampled at 2048Hz. During preprocessing, all 50Hz frequencies were filtered out to remove line noise. Then, epochs were created in a period from −500ms until 2750ms around stimulus presentation. Data were then re-referenced offline to the average of all electrodes. Then, channels with a noisy signal were removed and interpolated. On average, this was the case for 2.10 electrodes (SD=1.65, range=0-6). Afterward, a manual scan was conducted to delete epochs with clear (non-ocular) artifacts (e.g. sudden voltage jumps). Next, an independent component analysis (ICA) was performed on the data, and components reflecting eye movements or eye blinks were removed from the data. Then, the data were baselined to the time window −500 to -100ms prior to the stimulus presentation. Then, one last manual check was done to ensure that no epochs with artifacts were left. After preprocessing, the average number of epochs retained in the analysis was 676.6 (SD=53, range 479-718). Finally, EEG data were downsampled to 512Hz. These data were then also aligned to the time of the response (using a time window from - 200ms until 800ms) and to the time of the confidence response (using a time window from - 800ms until 100ms), in both cases keeping the same pre-stimulus baseline (i.e. following Feuerriegel et al., 2022). The EEG data was preprocessed using the MNE package in Python (Gramfort et al., 2013). The main findings reported in the manuscript were fully replicated when preprocessing was repeated using an alternative pipeline that applied average referencing after interpolating bad electrodes, omitted notch filtering, and included automatic rejection of trials exceeding ±150 µV on a predefined set of electrodes (Fz, F1, F2, CPz, Pz, POz, P1, P2).

### EEG analysis

Grand-average EEG waveforms were computed for all three epochs (stimulus-locked, response-locked, confidence-locked) for a set of a priori selected centro-parietal electrodes (CPz, Pz, Poz, P1 and P2) and for a set of fronto-central electrodes (Fz, F1 and F2). Differences between conditions (i.e. high vs. low confidence and positive vs. negative prior belief) were assessed using cluster-based permutation tests with an alpha value of .05 both for the cluster-forming phase and for comparing clusters against a permutation distribution of 1024 samples. Note that in this first step we decided to compare high vs low confidence, instead of showing the ERP separately per level of confidence, because data were too unbalanced to compute a stable ERP for each confidence level (for completeness, ERPs separately per level of confidence can be found in Supplementary Figure S1). As a follow-up, single-trial amplitudes for the centro-parietal and fronto-central electrode groups were extracted for the CPP, Pe and confidence-locked signal in a 200ms time window. For the CPP this was done around the empirically observed peak at 540ms post-stimulus (i.e. a window between 440ms and 640ms), for the Pe this was done in a time-window between 300ms and 500ms post-response (as suggested by Desender et al., 2021, p.9), for the confidence-locked signal this was done between −500 and -300 before the confidence response. Note that these time windows were chosen entirely blind to any potential effects of prior beliefs, thus avoiding any circularity regarding potential prior beliefs effects. In order to satisfy mixed model assumptions, these signals were z-scored separately per participant. These standardized single-trial voltages were then submitted to a linear mixed effects model with confidence, prior beliefs and their interactions as predictors, following the same approach for random slopes as for the behavioral analyses. Unlike ERP analyses, linear mixed-effects models are well suited for handling unbalanced data, as they can pool information across participants while accounting for both fixed and random effects.

We performed time-frequency analysis using a Morlet wavelet transformation to estimate power spectral density for frequencies ranging between 3 Hz and 30 Hz. The number of cycles used for wavelet transformation increased linearly from 1.5 cycles at the lowest frequency to 5 cycles at the highest frequency. This scaling of cycles ensures better frequency resolution at lower frequencies and better temporal resolution at higher frequencies. The time-frequency decomposition was done separately for each participant and each electrode within the set of centro-parietal or fronto-central electrodes. We computed percentage change in power between high and low confidence or between the positive and negative prior belief conditions, normalized by their sum to control for overall power differences. Afterwards, we averaged across electrodes within each ROI and applied a Gaussian smoother with a standard deviation of 2 (i.e. across the time and frequency dimension) to smooth the resulting decomposition. Note that the same results were obtained without applying a Gaussian smoother. Significant differences in power were assessed using cluster-based permutation testing at an alpha level of .05.

Finally, we carried out a set of multivariate decoding analyses on confidence and prior beliefs. Importantly, because there is an association between confidence and prior beliefs in the behavioral data, decoding analyses were carried out on a sub-selection of trials that were matched for both prior beliefs and confidence. Specifically, for each level of confidence in the positive prior beliefs condition, we randomly selected a trial in the negative prior beliefs condition with the same level of confidence. This process continued until no further matches were possible. Subsequently, prior beliefs (positive or negative) were decoded using a linear logistic regression decoder. This pipeline was then employed using a temporal generalization approach where classifiers were trained and tested on each timepoint (King & Dehaene, 2014), using five-fold cross-validation. Classification performance was evaluated using ROC AUC. Confidence was decoded as a continuous variable using ridge regression. Model performance was assessed using cross-validation with Spearman correlation as the scoring metric. To assess whether the classifier trained to decode continuous confidence was able to predict prior beliefs, within each fold the classifier was trained to predict continuous confidence (as described above), and we then computed the average (continuous) prediction for the held-out data separately for positive and negative prior beliefs. This difference score was subsequently used as a scoring metric. For all decoding analyses, above-chance decoding performance was assessed using cluster-based permutation testing at an alpha level of .05.

## Results

### Prior beliefs causally influence decision confidence

We analyzed the behavioral data using (generalized) linear mixed effects models with the dependent variables being either accuracy, reaction times or confidence, and as fixed effects structure the variables difficulty (3 levels: easy, medium or hard), manipulation type (2 levels: feedback or training difficulty), prior beliefs (2 levels: positive or negative) and their interaction. Behavioral results are shown in Figure 1C. For accuracy, there was only a main effect of trial difficulty, X²(2) = 1149.75, p < .001, indicating that accuracy increased as the task was easier (easy trials: M = 93.0%, medium trials: M = 84.5%, hard trials: M = 69.8%). There were no other significant effects, ps > .069, BFs < .54. For reaction times on correct trials, there was a main effect of trial difficulty, F(2,28.5) = 37.82, p < .001, indicating that RTs became shorter as the task was easier (easy trials: M = .891s, medium trials: M = .962s, hard trials: M = 1.018s). There was an unexpected interaction between manipulation type and prior beliefs, F(1,16321.5) = 11.87, p < .001. Post-hoc comparisons revealed that RTs were faster when participants received positive (M = .913s) versus negative (M = .964s) comparative feedback, z = 2.88, p = .004, whereas all other comparisons were not significantly different, ps > .148. No other effects were significant, ps > .064, BFs < .25.

Most importantly, we then examined the influence of our manipulation on confidence. There was a main effect of trial difficulty, F(2,19991) = 517.88, p < .001, indicating that confidence increased as the task was easier (easy trials: M = 4.12, medium trials: M = 3.85, hard trials: M = 3.66). Critically, there was a main effect of prior beliefs, F(1,20006)= 553.81, p < .001, with participants expressing higher confidence after receiving positive comparative feedback or training on an easy task (M = 4.01) compared to negative comparative feedback or a difficult training (M = 3.73). Although there was an interaction between prior beliefs and manipulation type, F(1,20003) = 29.43, p < .001, the prior beliefs effect was highly significant and robust in both the comparative feedback condition (M_positive-negative_ = .35, z = 20.41, p < .001) and the training difficulty condition (M_positive-negative_ = .22, z = 12.84, p < .001). All other effects were not significant, ps>.052. Note that the effect of feedback on confidence was much stronger compared to its effect on RTs, ruling out that this reflects a downstream effect. Indeed, when adding RTs to the model as an additional single-trial predictor, the main effect of feedback condition remained highly significant, F(1,2000) = 506.16, p < .001.

### Computational account of prior beliefs

In previous work, we have shown that the influence of prior beliefs on the computation of decision confidence can be accounted for within the drift diffusion model by incorporating an additional parameter, termed subjective drift, which captures the prior beliefs of the participant (Van Marcke et al., 2024). Specifically, confidence within this model is quantified as the probability of a choice being correct, conditional on the accumulated evidence and the time spent accumulating. Critically, this probability depends on the ability to perform the task: all else being equal, the probability of being correct is higher in easy compared to difficult tasks; so it is optimal for an agent to report higher confidence when they (merely) hold the prior belief that the task is easy (versus difficult). The subjective drift rate parameter controls the probability map used to map a certain level of evidence and reaction time onto confidence (i.e. the shape of the heatmap in Figure 1B). The subjective drift rate parameter is on the same scale as the regular drift rate, and thus captures prior beliefs about task performance in that it can be thought of as one’s subjective estimation of their regular drift rate. When fitting this model to the current data, we thus expected the model to capture the finding that the prior beliefs manipulation affects confidence, and it should do so by estimating a higher subjective drift rate in the positive condition. Indeed, the subjective drift rate was significantly affected by the prior beliefs condition, F(1,29) = 11.53, p = .002. We found no interaction between the prior belief condition and manipulation type, F(1,268) = 3.38, p = .067, further confirming that both manipulation types influenced confidence in a similar manner. Thus, this result confirms that manipulating prior beliefs, either via manipulative comparative feedback or via differences in training difficulty, causally influenced decision confidence via the same underlying mechanism: by changing the expected probability of being correct (represented by the subjective drift rate).

Even though no main effect of prior belief conditions was observed on accuracy and RT (i.e. during the behavioral analysis reported earlier), drift rate and non-decision time were slightly modulated by prior beliefs condition, drift rate: F(1,30) = 4.67, p = .039; non-decision time: F(1,29) = 4.33, p = .046. There was an interaction with manipulation type for drift rates, F(1,272) = 5.89, p = .016, but not for non-decision time, F(1, 1269) = 2.99, p = .085. Post-hoc comparisons revealed that drift rates were slightly higher after receiving positive compared to negative feedback, t(86) = -3.14, p = .002, whereas no difference was found in the training difficulty manipulation, t(84) = -.10, p = .923. This latter result is consistent with the slower RTs observed in the negative feedback condition compared to the positive feedback condition. Lastly, the boundary was not modulated as a function of prior beliefs condition, F(1,29) = .17, p = .682. Even though there was a significant prior belief condition X manipulation type interaction, F(1,267) = 4.94, p = .027, there was no effect of the prior belief condition on boundary in any manipulation type, ps > .406. As can be seen in Figure 1C, the computational model closely captured the behavioral findings.

Given that both feedback manipulation types influenced confidence in a highly similar manner, in the remainder of the manuscript we will ignore the specific manipulation type and only consider prior beliefs (positive/negative). By doing so, we ensure that we have sufficient experimental power for the EEG analysis and that the effects of prior belief in the EEG data are domain-general (i.e. regardless of the specific implementation of the prior belief).

### Parietal neural signals track accumulated evidence, not confidence

In the previous section, we established that our prior beliefs manipulation effectively influenced the reported level of decision confidence, and it does so by changing the mapping between the accumulated level of evidence and the probability of being correct. Next, we turned towards the EEG data to test the resulting prediction that common neural choice signals reflect accumulated evidence but not confidence.

Figure 2A shows the grand-average ERP of a set of centro-parietal electrodes (CPz, Pz, POz, P1, and P2) separately for trials rated with high confidence (‘certainly correct’ or ‘probably correct’, i.e., 4 or 5) or low confidence (‘certainly or probably wrong’, ‘guess wrong’, or ‘guess correct’, i.e., 1, 2, or 3). A cluster-based permutation test identified a clear modulation of the stimulus-locked CPP component by confidence, in a time window ranging between 440ms until the end of the epoch at 700ms, p = .002. The same modulation of the CPP by confidence was also seen when locking the data to the response, with a significant cluster ranging between the start of the epoch at -100ms before the response until 40ms after the response, p = .030. More importantly, there was a clear modulation of the response-locked Pe component by confidence in a time window ranging between 190ms and 660ms post-response, p = .005. Finally, when locking the data to the confidence response there were no significant clusters, ps > .085. In sum, the grand-average ERPs replicate the previously reported findings that both the centro-parietal CPP component (Desender et al., 2019; Dou et al., 2024; Feuerriegel et al., 2022; Gherman & Philiastides, 2018; Grogan et al., 2023) and the centro-parietal Pe component (Boldt & Yeung, 2015; Desender et al., 2019; but see Feuerriegel et al., 2022; Grogan et al., 2023; Ko et al., 2024) are modulated by decision confidence.

**Figure 2.**
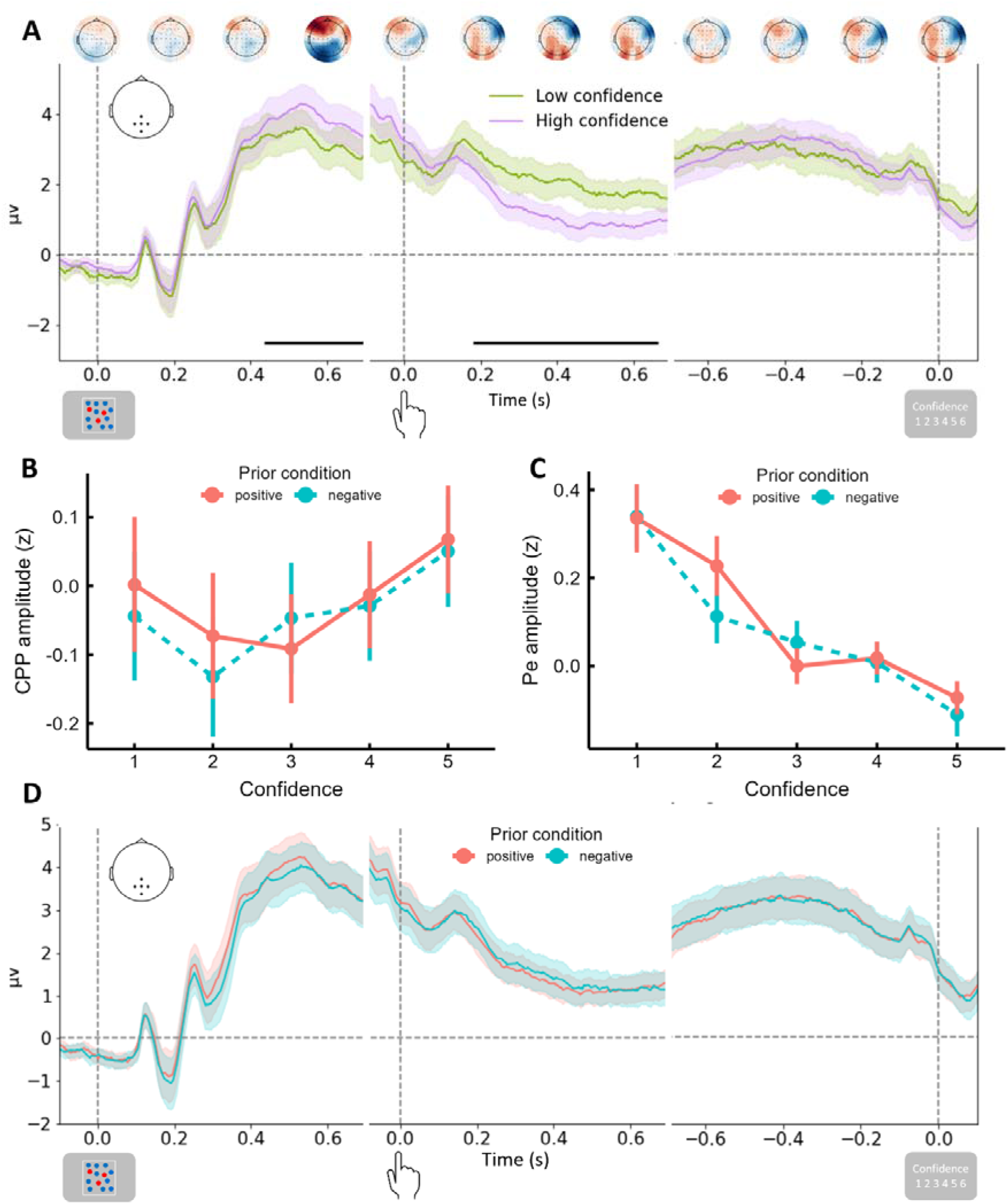
Parietal neural choice signals track accumulated evidence but not confidence. **A.** Cluster-based analysis of the centro-parietal EEG activity shows that both the stimulus-locked CPP and the response-locked Pe component were modulated as function of confidence (high vs low). Topographies show the difference in mean voltage between low and high confidence. **B-C.** The findings in A are compatible with these signals directly reflecting confidence and these signals reflecting accumulated evidence. To adjudicate between both, we tested whether these signals are sensitive to the influence of prior beliefs on confidence. Mixed-models showed that both CPP and Pe scale with the level of confidence, but are insensitive to prior beliefs, consistent with these signals reflecting the accumulated evidence. D. Cluster-based analysis of centro-parietal EEG activity did not show any difference between both prior beliefs conditions. Note: colored lines show group averages, and shaded bars reflect the standard error of the mean across participants. Horizontal black lines indicate significant time-windows as assessed by cluster-based permutation tests.

The finding that the CPP and PE closely scale with decision confidence (replicating previous work) is compatible with two distinct hypotheses: either these signals directly reflect confidence, or alternatively they reflect accumulated evidence which is used as input to compute confidence conditional on prior beliefs. To dissociate between both hypotheses, we investigated whether these signals are sensitive to the influence of prior beliefs on confidence. We extracted the average EEG amplitude from these centro-parietal electrodes in a 200ms window around the peak of the CPP (i.e. at 540ms) or a 200ms window ranging from 300ms to 500ms post-response. We then submitted these average EEG amplitudes to a mixed model using confidence (5 levels) and prior beliefs (2 levels: positive or negative) and their interaction as fixed effects.

For the CPP, we observed a main effect of confidence, F(4,19719) = 11.76, p < .001, with significant differences in CPP amplitude between adjacent confidence levels 3 and 4, z = 2.71, p = .007, and levels 4 and 5, z = 4.55, p < .001, but not between levels 1 and 2, p = .112, and between levels 2 and 3, p = .328, suggesting that the CPP is only sensitive to variations in confidence about subjectively correct trials (i.e. confidence reports ranging between guess correct and certainly correct) but not subjectively incorrect trials (i.e. confidence reports ranging between guess error and certainly error). Critically, there was no main effect of the prior belief condition, p = .511, BF = .06, nor was there an interaction between confidence and prior belief, p = .309, BF < .001 (see Figure 2B). To rule out that the effect of the prior belief was suppressed by confidence (because prior beliefs affect confidence), we reran the model including only prior beliefs condition as a predictor, which confirmed that prior beliefs did not affect average CPP amplitude, p = .376, BF < .001. Finally, the same key findings (main effect of confidence but not of prior beliefs) were observed when instead quantifying the CPP locked to the response (see Supplementary Figure S2).

For the Pe component, we observed a main effect of confidence, F(4,18635) = 28.30, p < .001, with significant differences in Pe amplitude between adjacent confidence levels 1 and 2, z = 2.80, p = .005, between level 2 and 3, z = 3.67, p < .001, and between levels 4 and 5, z = 5.53, p < .001, but not between levels 3 and 4, p = .491. Critically, there was no main effect of the prior belief condition, p = .128, BF = .07, nor an interaction between confidence and prior belief, p = .114, BF = .004 (see Figure 2B). Again, a model including only prior beliefs condition as a predictor showed no significant effect, p = .663, BF < .001.

To further demonstrate that centro-parietal signals do not reflect prior beliefs, we performed an additional cluster-based permutation test on the full-time course contrasting positive versus negative prior beliefs (see Figure 2D). As expected, there were no significant clusters in the stimulus-locked period, ps > .456, nor in the response-locked period (no significant clusters in the cluster forming phase), nor in the confidence-locked period (no significant clusters in the cluster forming phase). The same null effects were observed when separately analyzing the comparative feedback conditions (ps > .524 for stimulus-locked, no significant clusters for response-locked and confidence-locked data) and for the training difficulty conditions (ps > .310 for stimulus-locked, no significant clusters for response-locked and confidence-locked data). Finally, also when performing cluster-based analyses contrasting positive versus negative prior beliefs on individual electrodes (rather than contrasting the average across the centro-parietal electrodes), no significant clusters were identified.

### Oscillatory alpha and beta power track accumulated evidence, not confidence

Having established that common neural choice signals reflect accumulated evidence, but not confidence, we next investigated whether prior beliefs could be identified in oscillatory activity. This is motivated by previous work showing that power differences in pre-stimulus alpha oscillations are associated with the reported level of confidence (Samaha et al., 2017; Wostmann et al., 2018), and work showing that pre-stimulus oscillatory activity reflects prior expectations about upcoming responses (Lange et al., 2013). When looking at centro-parietal electrodes, we identified a clear pre-stimulus cluster (around −500ms until - 200ms) which dissociated between high and low confidence, p = .008 (see Figure 3A). The cluster was mostly pronounced at alpha power, but extended into the beta range, and reflected that lower alpha (and beta) power at baseline was associated with higher confidence about upcoming stimuli. In the response-locked data there were no significant clusters (ps > .085), but there was another significant cluster in the confidence-locked data (around −550ms until -300ms before the confidence response) dissociating high from low confidence which was mostly expressed within the beta power range, p = .043. Importantly, the same set of analyses contrasting positive versus negative prior beliefs did not reveal any significant clusters in neither stimulus-locked, ps > .959, response-locked, ps > .530, nor in the confidence-locked data, ps > .506 (see Figure 3B). Thus, although we were able to replicate previous work showing an association between oscillatory activity and confidence, we did not find any association between prior beliefs and oscillatory activity. The same set of analyses on the fronto-central set of electrodes revealed a similar pre-stimulus alpha effect on confidence, p = .012, but no other clusters for confidence, p > .064, and no significant clusters for prior beliefs, ps > .150.

**Figure 3.**
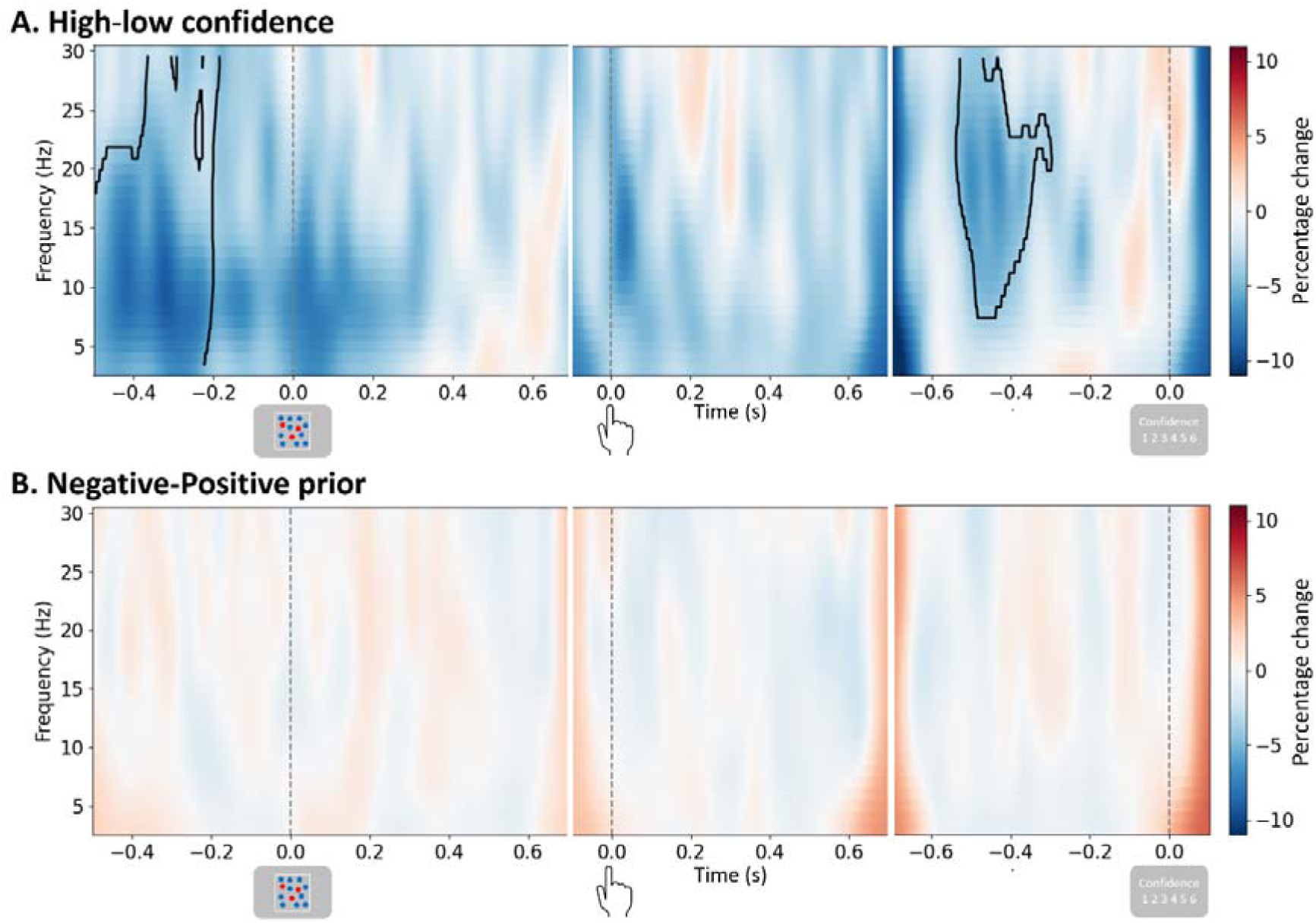
Oscillatory alpha and beta power track accumulated evidence, not confidence. **A.** Time-frequency decomposition of centro-parietal electrodes reveals two significant clusters sensitive to the difference between high and low confidence. A pre-stimulus cluster shows that reduced alpha (and beta) power is associated with higher confidence judgments, a cluster occurring before the confidence judgment shows that reduced beta power is associated with higher confidence. **B.** The same set of analyses contrasting trials where participants have positive versus negative prior beliefs does not reveal any significant clusters. Note: black lines indicate significant time-windows as assessed by cluster-based permutation tests.

### Decoding neural signatures representing prior beliefs and confidence

Having established that centro-parietal EEG signals and centro-parietal oscillatory activity do not reflect prior beliefs, we next attempted to unravel whether there are any neural signals in the EEG time course that do represent prior beliefs, and whether such signals are also sensitive to confidence. To this end, we turned towards multivariate time-resolved decoding. Specifically, we used a temporal generalization approach (King & Dehaene, 2014) and trained the classifiers on all timepoints across the three different events (stimulus-, response- and confidence-locked), and then testing performance of the classifiers on all timepoints across the three events. Importantly, because prior beliefs influence the level of confidence, it should be ruled out that the classifiers trained on one of these signals are actually picking up the other signal. Therefore, all decoding analyses were done on a subset of trials matched in terms of the frequencies of confidence and prior beliefs.

As a first sanity check, we tested whether it was possible to decode continuous confidence from EEG data using ridge regression. As shown in Figure 4A, there was a single large cluster of significant decoding starting around 400ms after the stimulus which extended into the entire response-locked and the entire confidence-locked epochs, p < .001. Decoding performance was reliably above chance level both on the diagonal as well as off-diagonal, suggesting a single stable pattern of neural activity that is consistent and sustained over time. This interpretation is confirmed by the scalp projections which show a reliably consistent pattern emerging around 400ms post-stimulus. Notably, although the scalp projections showed a clear centro-parietal locus, this extended more into frontal electrodes later in time.

**Figure 4.**
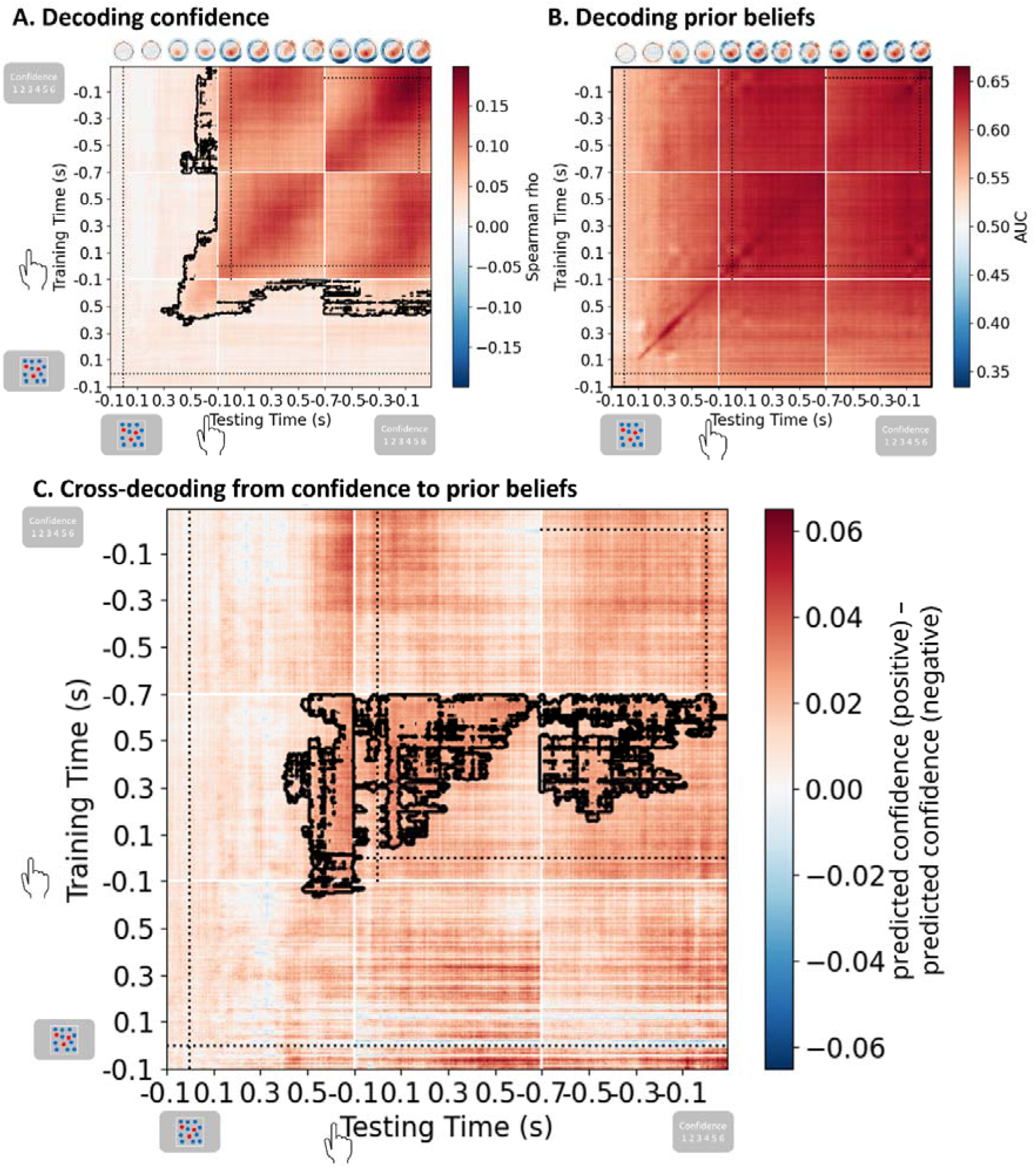
Cross-decoding prior beliefs and confidence using a temporal generalization approach across stimulus, response -and confidence-locked data. **A.** Decoding of continuous confidence levels using ridge regression. Starting from around 400ms post-stimulus it was possible to reliably decode confidence and this extended throughout the trial across response-locked and confidence-locked data both on -and off-diagonal, suggesting a single stable pattern of neural activity that is consistent and sustained over time. Scalp projections showing the decoder classifier weights identify a clear centro-parietal locus, extending more frontally later in the trial. **B.** Decoding of prior beliefs (positive vs. negative) revealed a single cluster visible in all three events both on -and off-diagonal. Starting slightly before the stimulus, the classifier was able to reliably decode the prior beliefs condition, and this extended throughout the trial. C. Cross-decoding performance of a classifier trained to predict continuous confidence and tested on its ability to predict prior beliefs. Cross-decoding was mostly possible when classifiers were trained on response-locked data, while during testing it was visible at the end of the stimulus-locked data, throughout the response-locked data and during part of the confidence-locked data. Note that the training for this analysis was similar to that of panel A, hence scalp projections are not included here to avoid duplication. Same conventions as in Figure 2.

Second, we tested whether we could reliably decode prior beliefs based on the EEG data. As shown in Figure 4B, there was a single large cluster of significant decoding which was seen in the entire stimulus-locked, the entire response-locked, and the entire confidence-locked epochs, p < .001. Similar to the decoding of confidence, there was reliable above chance level decoding both on the diagonal as well as off-diagonal, again showing that a single stable pattern of neural activity represents prior beliefs which is consistent and sustained over time. The scalp projections show a mostly centro-parietal locus for this effect.

Finally, we tested whether the neural signals associated with confidence were able to reliably predict the prior beliefs. To achieve this, we trained the classifier to predict continuous confidence based on the EEG data, and then for the held-out data we subtracted the predicted confidence values for the negative prior beliefs condition from the predicted confidence values for the positive prior beliefs condition. If this difference score is significantly above zero, it indicates that decoding of confidence carries relevant information about the prior beliefs. As shown in Figure 4C, there was consistent and reliable cross-decoding between confidence and prior beliefs, however the pattern was more scattered compared to the previous two analyses. The cluster-based permutation test identified a single significant cluster during which across-condition decoding was possible, p = .046. This cluster was most prominently visible in the response-locked data, yet already visible in the final timepoints of the stimulus-locked data. Interestingly, although the cluster did not extend into the confidence-locked data, there was off-diagonal decoding when classifiers were trained on the response-locked data but tested on the confidence-locked data. Such asymmetries in across-condition decoding often reflect a difference in signal-to-noise ratio (SNR) between the two conditions. Indeed, when restricting the analysis to the subset of participants who displayed numerically positive across-decoding in the response-locked data (i.e. participants with high SNR) we observed significant off-diagonal decoding, i.e. in all four quadrants obtained when combining response-locked and confidence-locked data.

### Frontal signals tracking both confidence and prior beliefs

In the previous sections, we established that centro-parietal signals are sensitive to the reported level of confidence but not to prior beliefs. Using multivariate decoding techniques, we found that there is clear and strong encoding of prior beliefs in EEG recordings, which even share some neural representations with decision confidence. Some of the scalp projections of the decoding seem to point to a frontal contribution to confidence and prior beliefs. Therefore, in a final set of analyses we examined whether there are frontal signals which represent both signals. Note that this set of analyses is not just inspired by our decoding results, but also by fMRI research showing that reported confidence has a frontal locus (Bang et al., 2020), and by recent EEG findings showing a frontal correlate of confidence (Lim et al., 2020; Feuerriegel et al., 2022).

Following Lim et al. (2020), we looked at the grand-average ERP of a set of frontal electrodes (Fz, F1, F2) separately for trials rated with high (4 or 5) or low (1, 2 or 3) confidence. Note that these are the exact same electrodes also tested in Lim et al., and thus the choice of electrodes was not inspired by the results of the decoding analyses. A cluster-based permutation test identified a clear modulation by confidence in a stimulus-locked time window ranging between 510ms until the end of the epoch at 700ms, p < .001, highly similar in timing to the CPP effect observed at centro-parietal electrodes (see Figure 5A). Different from the centro-parietal CPP component, however, a linear mixed model performed on this frontal signal with confidence, prior beliefs and their interaction as predictors showed main effects of both confidence, F(4,19598) = 4.04, p = .003, and of prior beliefs, F(1,76.5) = 6.02, p = .016, with no interaction between both factors, p = .146. As can be seen in Figure 5B, the frontal signal consistently reached a higher amplitude in the negative versus the positive prior beliefs condition. Follow-up tests for confidence showed differences between adjacent levels of confidence 1 and 2, z = 2.14, p = .033, between 3 and 4, z = 3.20, p = .001, but not between the other levels, ps > .662. One notable difference with the parietal CPP signal, is that the frontal signal does not show a monotonic association with confidence, but instead shows an inverted-U shape, being most negative for trials judged with certainty (i.e. either certainly wrong or certainly correct). However, to more conclusively rule out the possibility that the parietal and frontal signals simply reflect opposite poles of a single neural dipole, we performed two additional analyses. First, we ran the same model again but now included the centro-parietal CPP signal as an additional single-trial regressor in the model. Even after accounting for variation explained by the CPP, F(1,19925) = 2980.55, p < .001, there still were main effects of confidence, F(4,19361) = 2.96, p = .019, and prior beliefs, F(1,71.3) = 5.43, p = .022. Second, when first regressing out the centro-parietal CPP from the frontal signal and then submitting these residuals to the analysis, we again observed significant main effects of confidence, F(4,19391) = 3.75, p = .005, and prior beliefs, F(1,89.4) = 5.49, p = .021.

**Figure 5.**
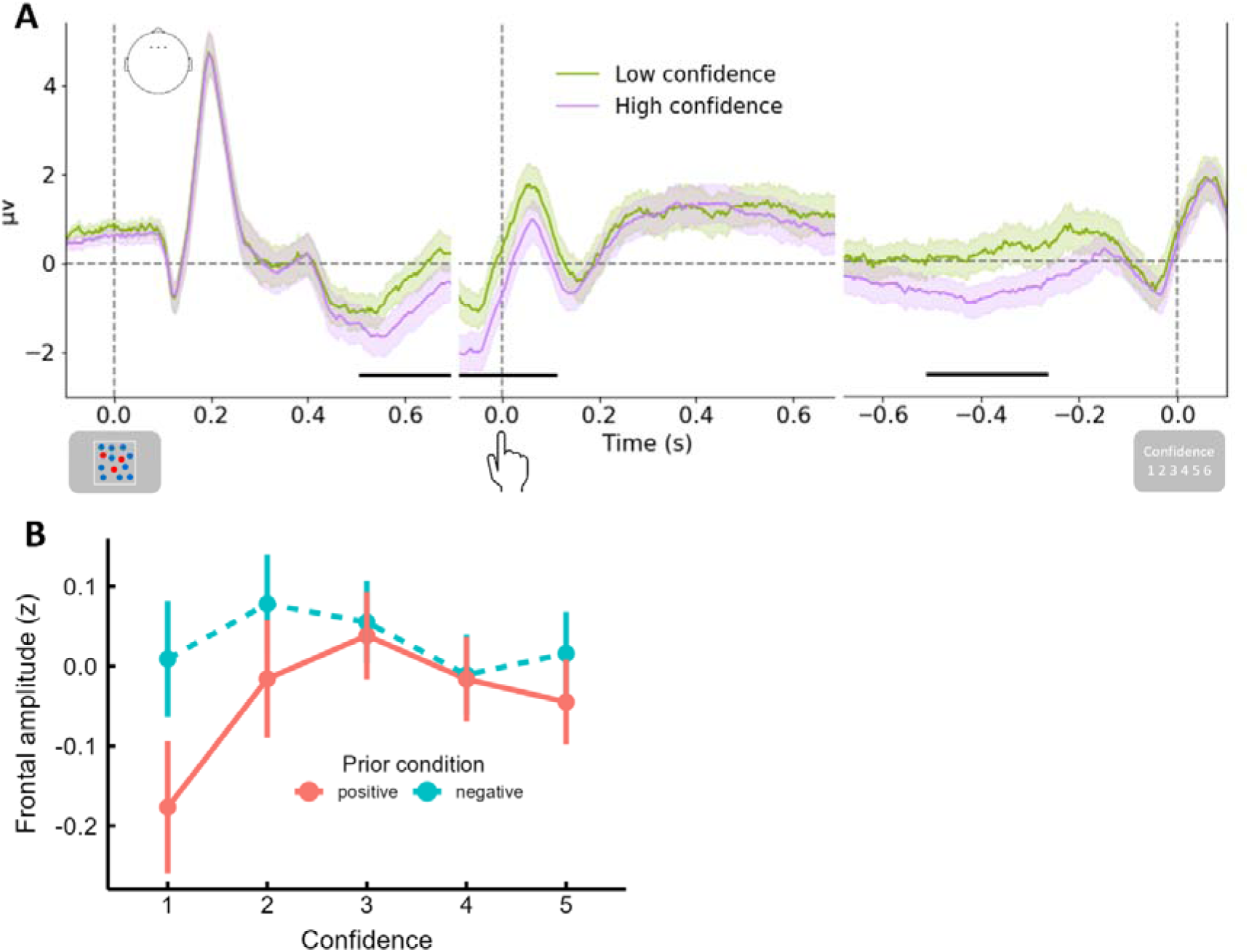
A frontal stimulus-locked signal encodes both prior beliefs and confidence. **A.** Cluster-based analysis of the frontal EEG activity shows both a stimulus-locked and confidence-locked signal that is modulated as function of confidence (high vs low). **B.** Follow-up analysis using mixed-models show that the stimulus-locked frontal signal scales with the level of confidence and with the prior beliefs condition, without an interaction between both signals.

Further inspection of the grand-average ERP revealed a similar effect of confidence on the frontal signal in the response-locked epochs between -200ms before and 110ms after the response, p = .005. Likewise, there was a clear effect in the confidence locked epoch from −510ms until -270ms before the confidence response, p=.007. It is very likely, however, that both these signals simply reflect the same frontal signal as seen in the stimulus-locked epoch. The median reaction time across the entire sample is 860ms, whereas the median confidence reaction time is 322ms, thus a signal appearing 510ms before the confidence response is consistent with it being a signal between 510 and 780ms after the stimulus and the same holds for the pre-response effect seen in the response-locked data. Nevertheless, a similar mixed effects model on the confidence-locked data showed a main effect of confidence, F(5,17971) = 8.04, p < .001, but no main effect of prior beliefs, p = .103, BF = .10, nor an interaction effect, p = .199, BF < .001.

## Discussion

An increasing number of studies attempts to unravel the temporal unfolding of neural signals reflecting decision confidence. An emerging consensus is that centro-parietal signals (either before or after the response) correlate with the level of decision confidence. Recent modeling work, however, makes a key prediction: common neural choice signals that are often associated with confidence actually reflect the accumulated evidence that is used as input to quantify confidence, instead of directly reflecting confidence. To test this novel prediction, in the current work we causally manipulated prior beliefs about the ability to perform a task, which effectively dissociates evidence from confidence. Results of an EEG experiment confirmed the hypothesis and showed that centro-parietal choice signals, specifically CPP and Pe, reflect the accumulated evidence that is used by the system to compute confidence, instead of directly reflecting confidence. A set of decoding analyses revealed that the brain does hold shared representations for prior beliefs and confidence, and we finally report a frontal signal that is sensitive to both confidence and prior beliefs, and thus identifies as a candidate signal to fulfill the role of a true confidence signal.

### Common neural choice signals reflect evidence

A wealth of previous EEG studies has shown close associations between (pre- or post-decision) centro-parietal signals and the level of decision confidence (Boldt & Yeung, 2015; Desender et al., 2019; Dou et al., 2024; Feuerriegel et al., 2022; Gherman & Philiastides, 2015, 2018; Kopcanova et al., 2023; Lim et al., 2020; Rausch et al., 2020; Stone et al., 2024). Importantly, in the current work we were able to replicate these findings: when setting aside the manipulation of prior beliefs, we did observe a close association between the (pre-decision) CPP and decision confidence and between the (post-decision) Pe and confidence. Although some concerns have been raised about whether the latter association reflects an artifact of using a pre-response baseline (Feuerriegel et al., 2022; Ko et al., 2024), we replicated the monotonic association between confidence and the Pe despite using the pre-stimulus baseline for the response aligned analyses (see also Boldt & Yeung, 2015; Desender et al., 2019). Also, although the interpretation of the stimulus-locked CPP might be misleading (i.e. because high confidence tends to be associated with shorter RTs and thus different CPP latencies; Feuerriegel et al., 2022), we fully replicated our main findings when quantifying the CPP in a response-locked time window instead.

Importantly, however, without considering the prior beliefs manipulation, the association between CPP/Pe and confidence is consistent with two potential explanations: these signals directly reflect the computation of confidence (as predicted by most confidence models) or alternatively these signals reflect accumulated evidence which then feeds into the computation of confidence (as proposed by the recent model by Van Marcke et al., 2024). Our EEG findings are consistent with the latter view: even though at the behavioral level we observed clear differences in confidence between the two prior beliefs conditions, both the CPP and the Pe were unaffected by this manipulation. According to our modeling framework, these signals thus track the amount of accumulated evidence which is subsequently used to compute the probability of being correct conditional on the prior beliefs. These findings were conceptually replicated in oscillatory activity, where reported confidence scaled with pre-stimulus alpha activity and pre-confidence report beta activity (Samaha et al., 2017; Wostmann et al., 2018), but there was no oscillatory activity that correlated with prior beliefs. Note that our finding, that CPP and Pe are insensitive to the manipulated prior belief about the ability to perform the task, does not implicate that these signals are not sensitive to any sort of prior beliefs. Indeed, recent work has shown modulations of the CPP by decision-relevant priors, such as predictive cues about the upcoming decisions (Kelly et al., 2021). Finally, one potential risk of our within-subjects design (i.e. each participant took part in all four prior beliefs conditions while performing a different perceptual task) is that the four tasks might slightly differ in RTs and accuracy, risking that EEG component latencies might be different between tasks. Although the four tasks were indeed not perfectly matched, the main findings reported in the current manuscript were replicated (except for the decoding of confidence in which no significant clusters were identified, likely due to a loss of power) when repeating the analysis on a subset of the data only including the orientation and shape task, for which there were no differences in RTs nor accuracy.

### A neural signal tracking confidence

Having demonstrated that common neural choice signals reflect evidence, an obvious next question is which neural signal reflects the computation of decision confidence. Although the CPP and Pe did not reflect the influence of prior beliefs on confidence, multivariate decoding analyses demonstrated that there was considerable neural activity predictive of prior beliefs, and some of this neural activity could cross-classify between confidence and prior beliefs, suggesting a common neural signal. Interestingly, the observation that prior beliefs could be identified with multivariate but not univariate analytical approaches suggests that prior beliefs are encoded in distributed spatial patterns of neural activity. This is consistent with the idea that multivariate analyses can capture information that may remain undetectable in traditional univariate approaches (Grootswagers et al., 2017). Although speculative, we identified a frontal component which appeared sensitive to both confidence and prior beliefs. This frontal signal reached a higher amplitude for decisions made with low confidence, and interestingly also a higher amplitude for negative versus positive prior beliefs. This pattern is consistent with a signal that integrates evidence and prior beliefs into a joint confidence judgment. If this frontal signal indeed reflects the computation of confidence, this has relevant implications with regard to the interpretation of the Pe component, which appears later in time than the frontal signal and thus would come too late to reflect a signal informing confidence. Although speculative, this finding aligns well with a recent finding that the CPP reflects both pre- and post-decision accumulation, whereas confidence RTs are too short to be explained as the result of the Pe (Grogan et al., 2025).

The possibility that there exists a frontal signal reflecting confidence is consistent with EEG work by Lim and colleagues (Lim et al., 2020). Note that these authors observed a positive monotonic association between the frontal component and confidence, which differs from the current findings. One potential difference is that in Lim et al. participants were not allowed to report having made an error, and thus it is unclear what participants reported when they caught themselves doing so. Moreover, several fMRI studies have mostly associated decision confidence with frontal regions (Bang et al., 2020; Boldt & Gilbert, 2022; Fleming et al., 2010, 2012; Miyamoto et al., 2018; Yokoyama et al., 2010). Our results seem closely related to an fMRI study by Bang and colleagues (2020) who showed that the lateral prefrontal pole is involved in transforming an internal confidence signal represented in perigenual ACC into a context-dependent public report. Likewise, a potential explanation for the current findings is that frontal regions compute confidence by incorporating evidence accumulated by parietal regions conditional on prior beliefs. This interpretation is different from Bang and colleagues, though, in which the context-dependent mapping by the prefrontal pole was a strategic operation (i.e. to match the confidence of another person). Therefore, it remains to be seen in future fMRI work whether the current findings can be similarly explained by a frontal region combining accumulated evidence and prior beliefs into a confidence signal. Importantly, however, given that the focus of the current work was not on this frontal signal, and its discovery should therefore be seen as exploratory, future work should further unravel the robustness of this signal as a neural representation of decision confidence.

## Conclusion

By manipulating prior beliefs about the ability to perform a task, we were able to show that confidence judgments are computed based on the accumulated evidence conditional on prior beliefs. Given that two centro-parietal EEG signals commonly referred to as the CPP (for pre-decision) and the Pe (for post-decision) were both insensitive to the influence of prior beliefs on confidence, we were able to show that these signals reflect accumulated evidence and not confidence. We instead identified a frontal signal which serves as a prime candidate to represent confidence.

## Acknowledgements

This research was supported by a grant by the Research Foundation Flanders, Belgium (FWO-Vlaanderen No. G0B0521N) awarded to K. Desender, and a grant by the Research Foundation Flanders, Belgium (FWO-Vlaanderen No. 11E6423N) awarded to H. Van Marcke. The authors would like to thank John Grogan and Redmond O’Connell for useful comments on an earlier draft.

## References

1. Bang, D., Ershadmanesh, S., Nili, H., & Fleming, S. M. (2020). Private–public mappings in human prefrontal cortex. ELife, 9, 1–25. 10.7554/eLife.56477

2. Bates, D. M., Maechler, M., Bolker, B., & Walker, S. (2015). Fitting Linear Mixed-Effects Models Using lme4. Journal of Statistical Software, 67(1), 1–48.

3. Boldt, A., & Gilbert, S. J. (2022). Partially Overlapping Neural Correlates of Metacognitive Monitoring and Metacognitive Control. Journal of Neuroscience, 42(17), 3622–3635. 10.1523/JNEUROSCI.1326-21.2022

4. Boldt, A., & Yeung, N. (2015). Shared Neural Markers of Decision Confidence and Error Detection. Journal of Neuroscience, 35(8), 3478–3484. 10.1523/JNEUROSCI.0797-14.2015

5. Desender, K., Murphy, P., Boldt, A., Verguts, T., & Yeung, N. (2019). A Postdecisional Neural Marker of Confidence Predicts Information-Seeking in Decision-Making. The Journal of Neuroscience, 39(17), 3309–3319.

6. Desender, K., Ridderinkhof, K. R., & Murphy, P. R. (2021). Understanding neural signals of post- decisional performance monitoring1: An integrative review. ELife, 1–21.

7. Dou, W., Arango, L. J. M., Castaneda, O. G., Arellano, L., Mcintyre, E., Yballa, C., & Samaha, J. (2024). Neural Signatures of Evidence Accumulation Encode Subjective Perceptual Confidence. Psychological Science. 10.1177/09567976241246561

8. Feuerriegel, D., Murphy, M., Konski, A., Mepani, V., Sun, J., Hester, R., & Bode, S. (2022). Electrophysiological correlates of confidence differ across correct and erroneous perceptual decisions. NeuroImage, 259 (November 2021), 119447. 10.1016/j.neuroimage.2022.119447

9. Fleming, S. M., Huijgen, J., & Dolan, R. J. (2012). Prefrontal contributions to metacognition in perceptual decision making. The Journal of Neuroscience, 32(18), 6117–6125. 10.1523/JNEUROSCI.6489-11.2012

10. Fleming, S. M., Weil, R. S., Nagy, Z., Dolan, R. J., & Rees, G. (2010). Relating introspective accuracy to individual differences in brain structure. Science, 329(5998), 1541–1543. 10.1126/science.1191883

11. Gherman, S., & Philiastides, M. G. (2015). Neural representations of confidence emerge from the process of decision formation during perceptual choices. NeuroImage, 106, 134–143. 10.1016/j.neuroimage.2014.11.036

12. Gherman, S., & Philiastides, M. G. (2018). Human VMPFC encodes early signatures of confidence in perceptual decisions. ELife, 7, 1–28. 10.7554/eLife.38293

13. Grogan, J. P., Rys, W., Kelly, S. P., & O’Connell, R. G. (2023). Confidence is predicted by pre- and post-choice decision signal dynamics. Imaging Neuroscience, 1(July 2023), 1–23. 10.1162/imag_a_00005

14. Grootswagers, T., Wardle, S. G., & Carlson, T. A. (2017). Decoding Dynamic Brain Patterns from Evoked Responses: A Tutorial on Multivariate Pattern Analysis Applied to Time Series Neuroimaging Data. Journal of Cognitive Neuroscience, 39(4), 677–697. 10.1162/jocn

15. Herregods, S., Le Denmat, P., & Desender, K. (2023). Modelling Speed-Accuracy Tradeoffs in the Stopping Rule for Confidence Judgments. BioRxiv, 1–58.

16. Kelly, S. P., Corbett, E. A., & O’Connell, R. G. (2021). Neurocomputational mechanisms of prior-informed perceptual decision-making in humans. Nature Human Behaviour, 5(4), 467–481. 10.1038/s41562-020-00967-9

17. Kelly, S. P., & O’Connell, R. G. (2013). Internal and external influences on the rate of sensory evidence accumulation in the human brain. The Journal of Neuroscience, 33(50), 19434– 19441. 10.1523/JNEUROSCI.3355-13.2013

18. Kiani, R., Corthell, L., & Shadlen, M. N. (2014). Choice Certainty Is Informed by Both Evidence and Decision Time. Neuron, 84(6), 1329–1342. 10.1016/j.neuron.2014.12.015

19. Kiani, R., & Shadlen, M. N. (2009). Representation of confidence associated with a decision by neurons in the parietal cortex. Science, 324(5928), 759–764. 10.1126/science.1169405

20. King, J.-R., & Dehaene, S. (2014). Characterizing the dynamics of mental representations: the temporal generalization method. Trends in Cognitive Sciences, 18(4), 203–210. 10.1016/j.tics.2014.01.002

21. Ko, Y. H., Zhou, A., Niessen, E., Stahl, J., Weiss, P. H., Hester, R., Bode, S., & Feuerriegel, D. (2024). Neural correlates of confidence during decision formation in a perceptual judgment task. Cortex, 3, 248–262.

22. Kopcanova, M., Ince, R. A. A., & Benwell, C. S. Y. (2023). Two distinct stimulus-locked EEG signatures reliably encode domain- general confidence during decision formation. 1–55.

23. Lange, F. P. De, Rahnev, D. a, Donner, T. H., & Lau, H. (2013). Prestimulus Oscillatory Activity over Motor Cortex Reflects Perceptual Expectations. The Journal of Neuroscience, 33(4), 1400–1410. 10.1523/JNEUROSCI.1094-12.2013

24. Lim, K., Wang, W., & Merfeld, D. M. (2020). Frontal scalp potentials foretell perceptual choice confidence. Journal of Neurophysiology, 123(4), 1566–1577. 10.1152/jn.00290.2019

25. Miyamoto, K., Setsuie, R., Osada, T., & Miyashita, Y. (2018). Reversible Silencing of the Frontopolar Cortex Selectively Impairs Metacognitive Judgment on Non-experience in Primates. Neuron, 980–989. 10.1016/j.neuron.2017.12.040

26. Moran, R., Teodorescu, A. R., & Usher, M. (2015). Post choice information integration as a causal determinant of confidence: Novel data and a computational account. Cognitive Psychology, 78, 99–147. 10.1016/j.cogpsych.2015.01.002

27. Morey, R. D., & Rouder, J. N. (2014). BayesFactor: Computation of Bayes factors for common design. Retrieved from Http://Cran.r-Project.Org/Package=BayesFactor.

28. Mullen, K. M., Ardia, D., Gil, D. L., Windover, D., & Cline, J. (2011). DEoptim: An R package for global optimization by differential evolution. Journal of Statistical Software, 40(6), 1–26. 10.18637/jss.v040.i06

29. O’Connell, R. G., Dockree, P. M., & Kelly, S. P. (2012). A supramodal accumulation-to-bound signal that determines perceptual decisions in humans. Nature Neuroscience, 15(12), 1729–1735. 10.1038/nn.3248

30. Pleskac, T. J., & Busemeyer, J. R. (2010). Two-stage dynamic signal detection: A theory of choice, decision time, and confidence. Psychological Review, 117(3), 864–901. 10.1037/a0022399

31. Ratcliff, R., & McKoon, G. (2008). The Diffusion Decision Model1: Theory and Data for Two- Choice Decision Tasks. Neural Computation, 20, 873–922.

32. Rausch, M., Zehetleitner, M., Steinhauser, M., & Maier, M. E. (2020). Cognitive modelling reveals distinct electrophysiological markers of decision confidence and error monitoring. NeuroImage, 218(January), 116963. 10.1016/j.neuroimage.2020.116963

33. Samaha, J., Iemi, L., & Postle, B. R. (2017). Prestimulus alpha-band power biases visual discrimination confidence, but not accuracy. Consciousness and Cognition, 54, 47–55. 10.1016/j.concog.2017.02.005

34. Stone, C., Mattingley, J. B., Bode, S., & Rangelov, D. (2024). Distinct neural markers of evidence accumulation index metacognitive processing before and after simple visual. Cerebral Cortex, May.

35. Tuerlinckx, F., Maris, E., Ratcliff, R., & De Boeck, P. (2001). A comparison of four methods for simulating the diffusion process. Behavior Research Methods, Instruments, and Computers, 33(4), 443–456. 10.3758/BF03195402

36. Van Marcke, H., Le Denmat, P., Verguts, T., & Desender, K. (2024). Manipulating prior beliefs causally induces under- and overconfidence. Psychological Science, 1–18. 10.1177/09567976241231572

37. Vandekerckhove, J., & Tuerlinckx, F. (2007). Fitting the Ratcliff diffusion model to experimental data. Psychonomic Bulletin & Review, 14(6), 1011–1026.

38. Wostmann, M., Waschke, L., & Obleser, J. (2018). Prestimulus neural alpha power predicts confidence in discriminating identical auditory stimuli. European Journal of Neuroscience, 136(1), 23–42.

39. Yokoyama, O., Miura, N., Watanabe, J., Takemoto, A., Uchida, S., Sugiura, M., Horie, K., Sato, S., Kawashima, R., & Nakamura, K. (2010). Right frontopolar cortex activity correlates with reliability of retrospective rating of confidence in short-term recognition memory performance. Neuroscience Research, 68(3), 199–206. 10.1016/j.neures.2010.07.2041

